# Sex allocation plasticity in response to resource and pollination availability in the annual plant *Brassica rapa* (Brassicaceae)

**DOI:** 10.1101/2022.12.30.522342

**Authors:** Nikolet Kostur, Susana M. Wadgymar

## Abstract

**Premise of research:** Hermaphroditic plants reproduce as females by maturing seeds from fertilized ovules and as males by fertilizing the ovules of other plants. Sex allocation theory predicts a trade-off between investment in male and female function. Thus, to maximize fitness, selection should favor plasticity in resource allocation among individuals or flowers of the same plant in response to environmental conditions. As female reproduction is typically more costly while male reproduction is mate-limited, we predict greater investment in female function when resources are plentiful and in male function when pollination is limited.

**Methodology:** We investigated plasticity in sex allocation in the rapid cycling lineage of the hermaphroditic mustard species, *Brassica rapa*, in response to resource availability (altered pot size) and the pollination environment (unpollinated or fully pollinated). We assess investment in male function (anther length) and female function (ovary length) in flower buds produced at the onset of reproduction and in buds produced approximately 15 days later. We also measured traits often correlated with increased allocation to female (plant size) and male (flower size) function.

**Pivotal Results:** Larger plants had longer anthers, longer ovaries, and larger flowers at the onset of reproduction, resulting in similar anther:ovary length ratios across plants of different sizes. Independent of plant-size, plants produced smaller anthers at the onset of reproduction in the low resource treatment and larger flowers over the course of reproduction in the pollen-absent treatment. Furthermore, larger plants produced increasingly longer ovaries over the course of reproduction compared to smaller plants.

**Conclusions:** Our findings underscore the influence of condition on changes in sex allocation and correlated traits over time. Furthermore, we provide some additional supporting evidence that resource availability and the pollination environment can influence sex allocation and contribute cautionary advice on effective methods for experimentally eliciting and measuring sex allocation plasticity.

## INTRODUCTION

Plants have evolved a diverse array of reproductive systems and strategies to maximize their reproductive assurance and success (Cardoso et al. 2018). For instance, approximately 90% of flowering plants produce bisexual flowers that house both male and female reproductive organs in the same floral unit, making hermaphroditism one of the most common sexual systems among angiosperms (Brunet 1992; Crowley et al. 2017; Christopher et al. 2019). Ovules are the structures within plants that contain the egg cell, and while they provide greater fitness value for investment of resources in female function, they are large, have limited mobility, and are resource-intensive to produce (Case and Ashman 2005; Strelin and Aizen 2018). In contrast, pollen, which contain sperm cells, require relatively few resources to produce, are smaller, and are more mobile (Cruden 2000; Rapp et al. 2013). In addition to the primary sexual characters of pollen and ovules, plants invest resources into secondary attractive and accessory characters that support these gametes and the fitness value of investment in male function (e.g., flower size, floral nectar, etc.). Hermaphroditic plants do not always invest resources equally into the two sexual functions because male and female functions vary in their resource requirements and reproductive assurance (Horovitz 1978).

The sex allocation of a plant describes its pattern of resource investment into male and female gamete production, as well as traits correlated with increasing male and female fitness (Charnov 1987). In 1877, Charles Darwin posited that increased investment in one sexual function would reduce the resources available for the other sexual function (Darwin 1877). This assumed trade-off formed the basis of sex allocation theory in plants (Jones and Ratterman 2009), which suggests that reproductive resources are limited and predicts that plants will allocate their resources between the sexes to maximize individual fitness (Primack and Kang 1989; Brunet 1992; Campbell 2000). Although total reproductive resources can be divided into categories (male, female, and associated structures), sex allocation represents the proportion of allocated resources and is a relative rather than absolute measure (Brunet 1992). Resource investment patterns between sexes can vary among individuals or among flowers within the same individual due to spatial or temporal differences in environmental conditions (Stehlik et al. 2008). Differences in the contexts in which the sexual functions provide reproductive assurance suggest that plants would benefit from a plastic sex allocation strategy in which they could adjust patterns of sex allocation in response to the biotic and abiotic environment (Rademaker and de Jong 2000; Zhao et al. 2008; Moore and Pannell 2011; Pellmyr et al. 2020).

Fitness gain curves describe the relationship between investment in a sexual function category and the male or female reproductive success (fitness) achieved (Charnov 1987; Brunet 1992; Brunet 1992). The shape of the gain curves is determined by intrinsic and extrinsic factors, including resource availability of shade, light, water, and soil (Bertin 2007; Li et al. 2019), as well as factors like disease status, herbivory, and quantity of pollen released by neighboring plants (Harder and Thomson 1989). While resources and pollinator service affect both male and female fitness, the shapes of gain curves determine whether one sex function benefits more than the other as the availability of a given factor increases (Brunet 1992; Emms 1993). For example, in a pollen abundant environment, the male fitness of animal pollinated plants is likely to decelerate with increased investment in male function: as resource investment in male function (i.e. pollen production) increases, the rate of male fitness gain decreases due to factors such as pollinators becoming saturated with pollen after visiting a plant (Lloyd and Bawa 1984). In contrast, under these conditions, female fitness (e.g., number of seeds produced) should accelerate with increased investment in female function (i.e. fruit and seed set) (Korpelainen 1998; Dorken and Mitchard 2008). As such, female fitness gains more than male fitness from resource input in this example. The quantitative differences in rates of fitness gain are the basis of conditional sex allocation theory (West 2009). If plasticity in sex allocation is shaped by selection as gain curves predict, we should be able to elicit predictable plastic responses in sex allocation and correlated traits under specific environmental conditions.

Here, we investigate plasticity in conditional sex allocation in the rapid cycling hermaphroditic mustard species, *Brassica rapa* (Brassicaceae), in response to two treatments that are each predicted to increase investment in male or female function. We ask the following questions: (1) Do plants in a resource-abundant environment invest proportionately more in female function than plants in a resource-limited environment? (2) Do plants in a pollination-absent environment invest proportionately more in male function than plants in a pollination-abundant environment? (3) Do plant size and flower size correlate with investment in male vs. female function?

Based on findings from previous studies (Austen and Weis 2014), we predicted high-resource plants to have relatively higher investment in female function than low-resource plants because female fitness gains more than male fitness from resource input (Korpelainen 1998). Specifically, we expected that the flowers produced by plants in the resource-abundant treatment would invest more in female function (low anther:ovary length ratio) in comparison to the flowers produced by plants in the resource-limited treatment, which would invest more in male function (high anther:ovary length ratio). We also predicted that a pollination treatment would elicit plasticity in sex allocation within individuals, with plants in the pollination-abundant treatment investing more in female function over time than plants in the pollination-absent treatment, which would invest more in male function over time. In all treatments, and in both early and later-produced flowers, we predicted greater investment in female function to correlate with larger plant size and greater investment in male function to correlate with larger flower size.

## MATERIALS AND METHODS

### Study species

*Brassica rapa* is a wild mustard species belonging to the Brassicaceae family. Native to Eurasia, *B. rapa* is an annual, self-incompatible hermaphroditic plant that is an obligate outcrosser. Its bisexual flowers mature acropetally (In succession from the base to the apex of the plant, Sarkissian et al. 2001) and flower bud production continues long after the first flowers open (Austen and Weis 2014). We used the base population for rapid cycling *B. rapa* (stock 1-001, University of Wisconsin-Madison), which retains genetic variation in phenotypic and life history traits (Williams and Hill 1986). The small size, short generation time of approximately 40-50 days, and self-incompatibility of this population make it well-suited for short-term pollination experiments conducted over the lifespan of a plant.

### Experimental design

We characterized sex allocation of sequentially produced flowers on 144 rapid cycling *B. rapa* plants. Seeds were cold stratified for 2 days, after which seedlings were planted on 29 January 2021 in a soil mixture consisting of 6.41 g of Osmocote 14-14-14 controlled-release fertilizer (ICL Specialty Fertilizers, Summerville, SC, USA) for every liter of Sun Gro Sunshine Mix #4 Aggregate Plus potting soil (Sun Gro Horticulture, Agawam, MA, USA).

Plants were grown in a Davidson College laboratory under Solar System 550 grow lights (California LightWorks, Canoga Park, CA, USA) set to a red:white:blue light setting of 10:16:10, which turned on everyday at 06:00 and turned off at 20:00. The laboratory temperature remained near 26.8 degrees celsius, with 10 percent humidity. Plants were watered as necessary using under-watering trays.

To manipulate resource availability, seeds were randomly planted in either large pots (10.2 W x 10.2 L x 8.9 H inches, 39 cm^3^ volume) or small pots (8.9 W x 8.9 L x 6.4 H inches, 14 cm^3^ volume, Greenhouse Megastore, Danville, IL, USA). Pots were distributed across 8 total trays, with big and small pots organized in an alternating, checkerboard pattern within each tray. We planted two seedlings per pot and randomly thinned one seedling from each pot that had multiple germinants six days after planting. Plants began flowering approximately 19 days after planting and flowered for about 15-20 days thereafter. Seven plants died over the course of the experiment, leaving a total sample size of n=137.

To assess the impact of the pollination environment on pollen and ovule production in *B. rapa*, we randomly assigned plants from both pot size treatments to either pollination-abundant or pollination-absent treatments. To minimize inadvertent pollen transfer among plants, pollination treatments were applied to all plants within a tray, with four trays of plants per pollination treatment. For the pollination-abundant treatment, all open flowers on all plants were hand-pollinated indiscriminately every 1-2 days using a feather. In the pollination-absent treatment, plants were individually and loosely wrapped in clear cellophane after reproduction began to prevent pollination from neighboring plants. We selected extreme pollination treatments to elicit the strongest possible plasticity in sex allocation.

### Data collection

To measure the allocation of resources towards male vs. female function, we collected a total of 4 flower buds per plant approximately 1 day before we expected them to open, when the buds were swollen and the yellow petal color was visible through the sepals. The first pair of flower buds to mature on each plant (hereafter referred to as the early flower buds) were collected to measure sex allocation at the onset of reproduction, with differences in sex allocation across treatments indicative of plasticity in response to resource availability. To measure plasticity in sex allocation that may occur over time among flowers on the same plant, we collected a second pair of mature flower buds 15 days after the first set of flower buds were collected (hereafter referred to as late flower buds). Growth rates and rates of flower production varied by treatment, making it challenging to select an appropriate date to collect the late flower buds. Consequently, we chose a late flower bud collection date within the flowering duration of all plants that would standardize the amount of time that plants experienced experimental treatments. During both of these collection periods, we used a caliper to measure the stem diameter just above the cotyledons and the diameters of the two flowers that were next to open immediately after early and late flower bud collections, which we expect to be correlated with investment towards female and male function, respectively. We stored the flower buds in labeled tubes containing 70% ethanol, which does not affect flower structure or size (Austen and Weis 2014).

We measured the length of all six anthers and the ovary (specifically, the valve) for each of the flower buds collected using a EZ4W1 dissecting microscope and Leica Software (Leica Camera AG, Wetzlar, Germany). We were unable to count the number of ovules within the ovary or to estimate the number of pollen grains each flower bud contained. However, anther length in *B. rapa* predicts pollen quantity in each flower (Pollen = –56,401 + 9417 × (Length), r = 0.88, n = 88 plants, per (Austen and Weis 2014). Additionally, gynoecium size predicts ovule number in *Arabidopsis thaliana*, which is also a member of the Brassicaceae family (Cucinotta et al. 2020). These studies suggest that the anther-ovary length ratio can be used as a proxy for pollen-ovule ratios.

### Data analysis

We assessed the influence of resource availability (pot size) on average anther length, average ovary length, average anther to average ovary length ratio, flower size, and stem diameter at the onset of reproduction (i.e. at the time that early flower buds were collected) via generalized linear mixed models using the nlme package (Pinheiro et al. 2014) in R (ver. 4.0.3; R Development Core Team, https://www.R-project.org). Average anther length was calculated as the average of all six anther lengths found in each flower bud. We included a fixed effect for pot size and included initial stem diameter as a covariate. Additionally, we included tray as a random term to account for any microhabitat differences that may have existed across the study space. A significant effect of pot size would indicate that the response variables were plastic in response to resource availability, independent of any variation that is explained by plant size at reproduction. We visually assessed residual normality and residual homogeneity across the continuous covariate. Variance heterogeneity between pot size treatments or across plant sizes were corrected using error variance covariates if they reduced AIC values relative to models without them (Zuur 2009).

We assessed the influence of resource availability and the pollination environment on changes in average ovary length, average anther length, average anther to average ovary length ratio, flower size, and stem diameter that occurred from early to late flower bud collection in models identical to those described above, but with the pollination treatment and its interaction with resource treatment included as additional fixed effects. A significant effect of resource or pollination treatment would indicate that response variables were plastic over the course of reproduction in response to variation in the resources or pollination they experience.

## RESULTS

### Sex allocation at the onset of reproduction

Larger plants had longer anthers, longer ovaries, and larger flowers at the onset of reproduction, resulting in similar anther:ovary length ratios across plants of different sizes (Table 1, Figure 1). Independent of plant size, average anther lengths were 6.2% longer in smaller pots (Figure 1a), although this difference was not large enough to impact the anther:ovary length ratio (Table 1). The data suggest that differences in condition do not affect patterns of sex allocation at the onset of reproduction. In addition, limited resources can increase investment in male function consistently across a range of plant conditions.

**Table 1.**
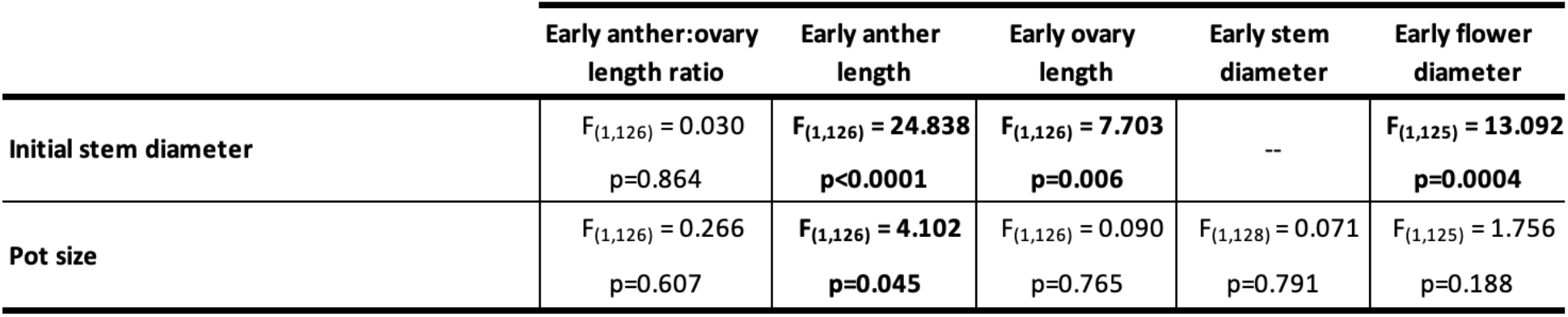
Effects of resource availability (pot size) and plant size on plasticity in sex allocation and correlated traits at the onset of reproduction. The pollination treatment was not included in these analyses because flower buds were collected prior to any flowers opening and being pollinated. Results were derived from generalized linear mixed models, with tray as a random effect. Boldfaced values are statistically significant at the p<0.05 level.

**Figure 1.**
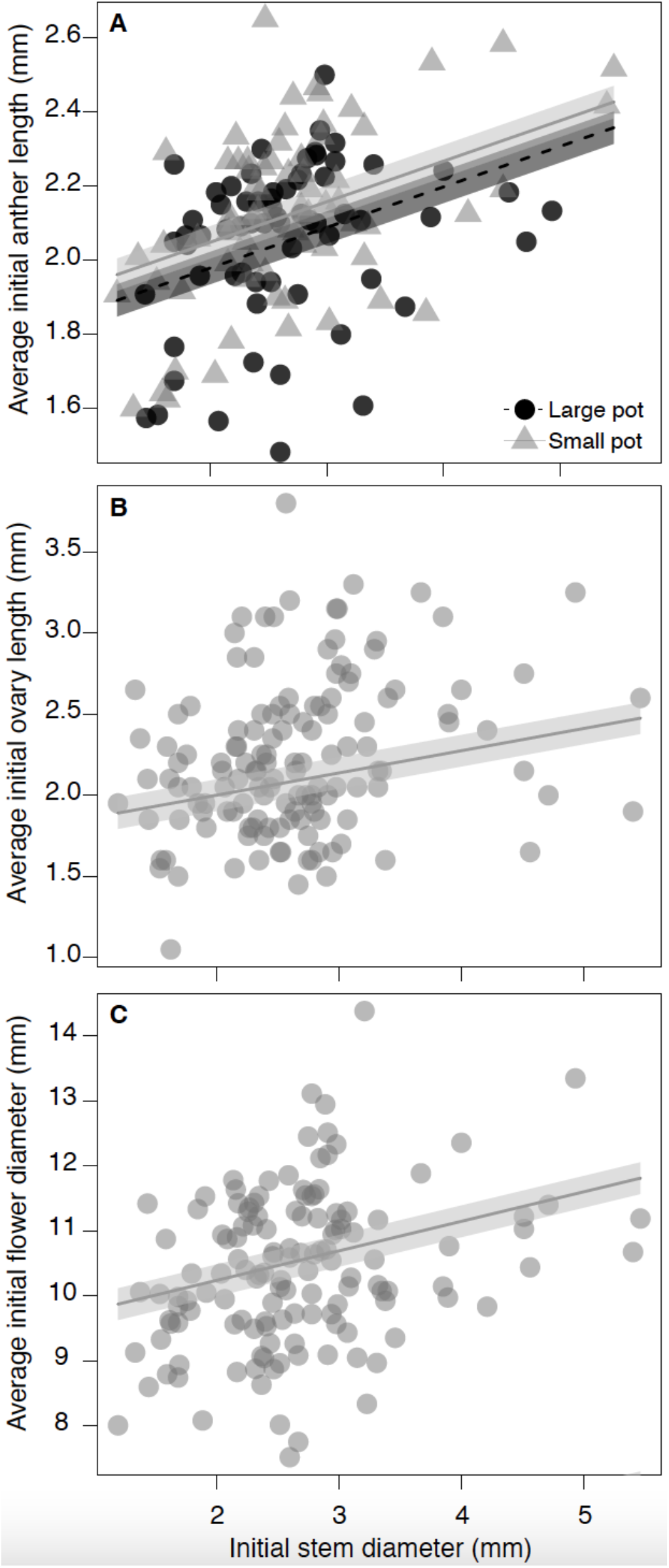
The associations between stem diameter at the onset of reproduction and (A) average initial anther length, (B) average initial ovary length, and (C) average initial flower diameter. Panel (A) also demonstrates the influence of resource availability (via differences in pot size) on average initial anther length. Trend lines and 95% confidence intervals were extracted from the results of generalized linear mixed models that included all other explanatory variables and incorporated tray as a random effect, as shown in Table 1.

### Sex allocation over the course of reproduction

Our pot size treatment influenced plant condition as we expected, with a marginally significant increase in stem diameter for plants in large pots (Table 2, Fig 2a). Plant size at the onset of reproduction did not explain variation in shifts of average anther or the anther:ovary length ratio over time, nor did pot size or pollination treatment influence the changes in average anther length, ovary length, or their ratio (Table 2). However, changes in the average ovary length did vary with plant size at the onset of reproduction, such that larger plantsproduced larger ovaries over time while the opposite pattern was observed for smaller plants (Table 2, Figure 2b). In addition, the pollination treatment did influence changes in flower size independent of plant size, with plants in the pollen-absent treatment producing larger flowers over time and plants in the pollen-abundant producing smaller flowers over time (Figure 2c).

**Table 2.** The effects of resource availability (pot size), pollination treatment, their interaction, and plant size on plasticity in sex allocation and correlated traits over the course of reproduction. Results were derived from generalized linear mixed models with tray as a random effect. Boldfaced values are statistically significant at the p<0.05 level.

|  | Changes in<br>anther:ovary<br>length over time | Changes in<br>Anther length<br>over time | Changes in in<br>ovary length<br>over time | Changes in<br>stem diameter<br>over time | Changes in<br>flower size<br>over time |
| --- | --- | --- | --- | --- | --- |
| **Initial stem diameter** | $F_{(1,118)} = 3.276$<br>$p = 0.073$ | $F_{(1,118)} = 2.351$<br>$p = 0.128$ | **$F_{(1,118)} = 6.525$**<br>**$p = 0.012$** | $F_{(1,122)} = 1.337$<br>$p = 0.250$ | $F_{(1,120)} = 2.211$<br>$p = 0.140$ |
| **Pot size** | $F_{(1,118)} = 1.529$<br>$p = 0.219$ | $F_{(1,118)} = 2.325$<br>$p = 0.130$ | $F_{(1,118)} = 0.450$<br>$p = 0.504$ | $F_{(1,122)} = 3.730$<br>$p = 0.056$ | $F_{(1,120)} = 1.946$<br>$p = 0.166$ |
| **Pollination treatment** | $F_{(1,6)} = 0.786$<br>$p = 0.410$ | $F_{(1,6)} = 0.721$<br>$p = 0.428$ | $F_{(1,6)} = 0.246$<br>$p = 0.637$ | $F_{(1,6)} = 2.125$<br>$p = 0.195$ | **$F_{(1,6)} = 8.761$**<br>**$p = 0.025$** |
| **Pot size x Pollination treatment** | $F_{(1,118)} = 0.014$<br>$p = 0.907$ | $F_{(1,116)} = 0.735$<br>$p = 0.393$ | $F_{(1,116)} = 0.009$<br>$p = 0.924$ | $F_{(1,121)} = 0.000$<br>$p = 0.994$ | $F_{(1,120)} = 0.855$<br>$p = 0.357$ |

**Figure 2.**
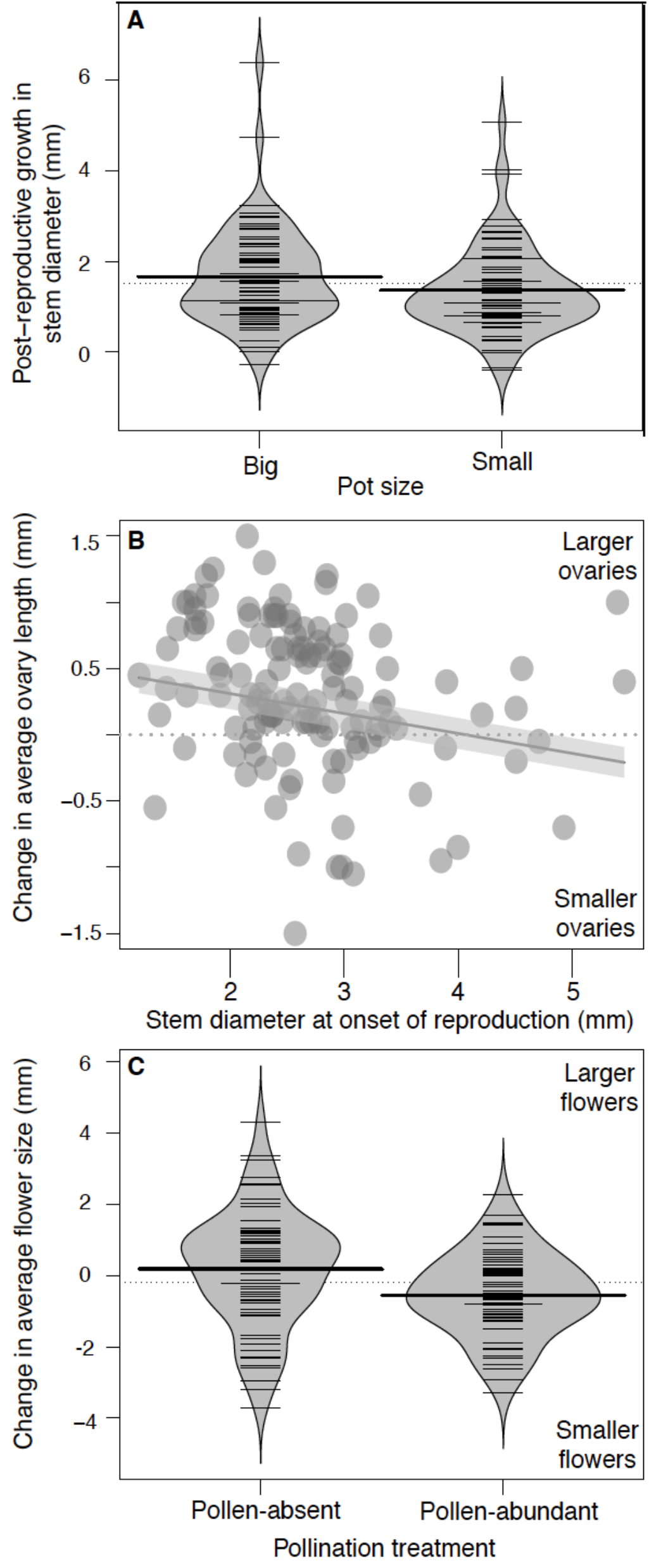
(A) Beanplots demonstrating the association between resource availability (pot size) and post-reproductive growth. Thin lines represent a one-dimensional scatter plot, thick lines represent the average anther length within each treatment, and the dotted horizontal line reflects the grand average including plants of both treatments. (B) The relationship between stem diameter at the onset of reproduction and changes in average ovary length over time. The horizontal dotted, grey line at y=0 reflects no plasticity in ovary length over time. The trend line and 95% confidence interval was drawn from the results of generalized linear mixed models that included all other explanatory variables and incorporated tray as a random effect, as shown in Table 2. (C) Beanplots demonstrating the association between the change in flower diameter between flower bud collection times and the pollination treatment. Thin lines represent a one-dimensional scatter plot, thick lines represent the average anther length within each treatment, and the dotted horizontal line at y=0 reflects no plasticity in flower size.

The effects we observed were in alignment with our predictions, suggesting that plants in resource-rich environments invest more in female function than they do in male function, thus becoming functionally more female over time. Furthermore, our data demonstrate that pollen-limited conditions elicit investment in larger flowers over time for plants of all sizes.

## DISCUSSION

We observed an influence of resource availability, plant size variation, and pollination environment on some aspects of sex allocation plasticity. Independent of plant-size, plants produced smaller anthers at the onset of reproduction in the low resource treatment (Figure 1a) and larger flowers over the course of reproduction in the pollen-absent treatment (Figure 2c). Furthermore, larger plants produced larger flowers at the onset of reproduction (Figure 1c) and increasingly longer ovaries over the course of reproduction (Figure 2b) compared to smaller plants. Our findings underscore the impacts that environmental conditions and resource status can have on sex allocation plasticity.

### Resource availability and plant size

In contrast to findings in past studies, the results of our experiment do not support our predictions that high-resource status (i.e. larger size or larger pots) promotes investment in female function over male function at the onset of reproduction or over the duration of flower production (size-dependent sex allocation theory, Shibata and Kudo 2016). In the self-incompatible hermaphrodite *Aconitum gymnandrum*, plant-size negatively correlated with initial sex allocation (measured as anther and gynoecium mass, Zhao et al. 2008). Similarly to our results, however, investment in female function increased over time in plants with greater access to resources (Zhao et al. 2008). These findings were obtained through a pot size and fertilizer manipulation treatment, in which plants were planted in smaller (14,726 cm^3^)/unfertilized pots and larger (28,274 cm^3^)/fertilized pots with a twofold and threefold increase in Nitrogen and Phosphorus, respectively. Similarly, plant size and sex allocation were affected by a pot size treatment in the self-incompatible hermaphroditic wild strawberry, *Fragaria virginiana*, in which half of plants were assigned to either small (75 cm^3^) or large (150 cm^3^) pots (Case and Ashman 2005). Plants in small pots were smaller compared to plants in the large pots two months after planting, and this size difference was maintained for the remainder of the experiment. As a result, ovule number per flower increased to a greater degree with increasing plant size, compared to pollen quantity per flower ((Case and Ashman 2005)), confirming that resource status can influence sex allocation.

Adjusting pot size is known to be a frequent and effective means for manipulating resources, with plants increasing biomass (measured in total dry mass) production by an average of 43% for every doubling of pot volume (Poorter et al. 2012). Although we detected a marginally significant effect of pot size on final plant size, we did not see significant effects of pot size on plant size at the onset of reproduction, suggesting that resource levels did not vary substantially among small and large pots during early ontogeny. We expect that we did not manipulate resource level severely enough and that plants in both treatments initially experienced comparable resource environments. In a pot size and fertilizer manipulation experiment on hermaphroditic individuals of *Fragaria virginiana*, plants were randomly assigned to either small (95 cm^3^) or large (280 cm^3^) pots, which differed in size to a greater extent than the 14 cm^3^ and 39 cm^3^ pots used in our study (Bishop et al. 2010). Nutrient levels were manipulated factorially with pot size, and both increased nutrients and larger pot size produced the largest plants at the end of the experiment (Bishop et al. 2010). We did not limit nutrients or resources such as light or water, which may explain the lack of effects of resource availability on plant size or anther:ovary length ratios.

Our chosen indicator of plant size may also explain the lack of influence of resource availability on initial plant size that we observed. Stem diameter is a common proxy for plant size in *Brassica rapa* (Franke et al. 2006; Hamann et al. 2018). However, it is possible that there are other, more reliable indicators of plant biomass that would have been appropriate to measure. For instance, taproot dry mass in *Brassica rapa* is positively correlated to height, shoot dry mass, and stem diameter (Austen and Weis 2014). In addition, correlations between root diameter and height varied among populations of *Brassica rapa* (0.68-0.93), suggesting that distinct measures of plant size do not perfectly covary (Mitchell-Olds 1996). Lastly, the rapid cycling life history of the *B. rapa* population we used may preclude stem diameter, or plant size in general, from being variable enough to address research questions related to size-dependent sex allocation.

### Pollination environment

There is ample evidence in the existing literature that pollen availability affects plasticity in sex allocation towards investment in the rarer gamete (Huang and Guo 2000; Rapp et al. 2013; Crowley et al. 2017; Pellmyr et al. 2020), with plants in pollen-absent environments investing more in male function and plants in pollen-abundant environments investing more in female function (Campbell 2000). For example, in a study on the effects of active insect pollination (insects intentionally collect pollen and deposit it on the stigma) versus passive pollination (stigma collects pollen as it contacts insects’ bodies) on pollen-ovule ratios in the hermaphroditic species *Yucca filamentosa, Hesperoyucca whipplei*, and *Lophocereus schottii*, actively pollinated plants (in which insects place pollen directly on the stigmas) had significantly lower pollen-ovule ratios than passively-pollinated plants of the same species (Pellmyr et al. 2020). These findings support predictions from sex allocation theory that plants in a pollen limited environment invest more in male function. Although we did observe larger flowers in our pollen-absent treatment, we observed no effects of pollination absence or abundance on sex allocation in our experiment.

The lack of effect of our pollination treatment on plasticity in sex allocation over time may be explained by our measurement of anther and ovary length as predictors of pollen and ovule production, limitations of the precision capabilities of the software we used to take measurements, and the timing of flower bud collection for the late pair of buds. In *Brassica rapa*, anther length has been shown to predict pollen number (Austen and Weis 2014) and gynoecium size can predict ovule number in *Arabidopsis thaliana*, another member of the Brassicaceae family (Cucinotta et al. 2020). We thus felt confident in using anther and ovary lengths as a proxy for pollen and ovule number, respectively. However, we did not confirm that ovary and anther lengths correlated with gamete number or size in the population of rapid cycling *B. rapa* that we used in our experiment. It’s possible that anther and ovary lengths are inappropriate measures of sex allocation in our study population. Furthermore, many plants continued to flower long after the late flower buds were collected, indicating that the collections for the late pair of flower buds might have occurred too early to detect any effects of the pollination treatments on plasticity in sex allocation. Klinkhamer and de Jong (2002) suggest that it is difficult to measure trade-offs between anther and ovary production in manipulation experiments, and instead suggest that researchers rely on genetic correlations to assess these trade-offs.

### Life history traits

Our hypotheses explored the influence of resource status and the pollination environment on sex allocation. While the timing of flowering was not a focal trait in our study, we did measure it and explore its association with our response variables (Appendix 1).

Independent of plant size at the onset of reproduction, we detected a non-linear association between flowering onset dates and sex allocation, such that early and late flowering plants were functionally more male than intermediate flowering plants. In addition, early flowering plants became functionally more female over time compared to intermediate and later flowering plants, where sex allocation remained the same over the duration of flowering. Including flowering onset as a covariate in our models didn’t change any of our results regarding the pot size and pollination treatments, but it did alter the significance of plant size on sex allocation in some cases (Appendix 1). These results are in agreement with those of Austen and Weis (2014), who found that initial sex allocation steadily increased in femaleness between early- and late-flowering plants, while anther-ovary ratios decreased (sex allocation became more male) over time in later flowering plants. Our data suggest that exploring how sex allocation relates to phenology and life history variation may be an intriguing future line of investigation.

## Conclusion

Our findings underscore the important influence of resource-status on patterns of sex allocation. Furthermore, we provide some additional supporting evidence that the pollination environment can influence sex allocation to secondary traits, and contribute cautionary advice on effective methods for experimentally eliciting and measuring sex allocation plasticity. We suggest that it would be worthwhile to monitor sex allocation in studies examining life history variation, particularly in the context of pollinator decline, shifting phenology, and decreases in the duration of growing seasons under climate change. Future work could also explore the influence of parental resource and pollination conditions on the sex allocation of their offspring (transgenerational plasticity) or could further explore the link between sex allocation and reproductive timing by using photoperiod manipulations to alter the timing of reproduction.

## ACKNOWLEDGEMENTS

We are grateful for funding support provided by Davidson College to S.M.W. and to members of the Wadgymar lab who assisted with pollination treatments, moral support, and breaking down and repeating this experiment due to the institutional shutdowns associated with COVID-19.

