## Appendix for "Sex allocation plasticity in response to resource and pollination availability in the annual plant *Brassica rapa* (Brassicaceae)": Appendix_Kostur and Wadgymar.pdf

**Summary of contents:** Our hypotheses explored the influence of resource status and the pollination environment on sex allocation. While the timing of flowering was not a focal trait in our study, we did measure it and explore its association with our response variables. We present those analyses below.

**Table 1.** Effects of resource availability, flowering time, and plant size on of plasticity in sex allocation and correlated traits at the onset of reproduction. The pollination treatment was not included in these analyses because flower buds were collected prior to any flowers opening and being pollinated. Results were derived from generalized linear mixed models, with tray as a random effect. Boldfaced values are statistically significant at the p<0.05 level.

| | Early anther:ovary length ratio | Early anther length | Early ovary length | Early stem diameter | Early flower diameter |
|---|---|---|---|---|---|
| **Days from germination to flowering (linear)** | **$F_{(1,124)}$ = 32.252 p<0.0001** | $F_{(1,125)}$ = 1.727 p=0.191 | **$F_{(1, 124)}$ = 17.922 p<0.0001** | **$F_{(1,126)}$ = 49.903 p<0.0001** | **$F_{(1,124)}$ = 6.014 P=0.016** |
| **Days from germination to flowering (quadratic)** | **$F_{(1,124)}$ = 22.331 p<0.0001** | -- | **$F_{(1,124)}$ = 20.213 p<0.0001** | **$F_{(1,126)}$ = 21.624 P<0.0001** | -- |
| **Initial stem diameter** | $F_{(1,124)}$ = 0.034 P=0.854 | **$F_{(1,125)}$ = 25.363 p<0.0001** | **$F_{(1,124)}$ = 9.556 p=0.0025** | -- | **$F_{(1,124)}$ = 13.460 P=0.0004** |
| **Resource availability** | $F_{(1,124)}$ = 0.882 p=0.249 | **$F_{(1,125)}$ = 4.848 p=0.030** | $F_{(1,124)}$ = 0.028 p=0.868 | $F_{(1,126)}$ = 0.446 p=0.505 | $F_{(1,124)}$ = 1.120 P=0.292 |

**Table 2.** The effects of resource availability, pollination treatment, flowering time, and plant size on plasticity in sex allocation and correlated traits over the course of reproduction. Results were derived from generalized linear mixed models with tray as a random effect. Boldfaced values are statistically significant at the p<0.05 level.

| | Changes in anther:ovary length over time | Changes in Anther length over time | Changes in in ovary length over time | Changes in stem diameter over time | Changes in flower size over time |
|---|---|---|---|---|---|
| **Days from germination to flowering (linear)** | **$F_{(1,116)}$ = 24.834 p<0.0001** | $F_{(1,116)}$ = 0.399 p=0.529 | **$F_{(1,117)}$ = 9.534 p=0.003** | $F_{(1,121)}$ = 0.157 p=0.692 | $F_{(1,119)}$ = 1.186 p=0.278 |
| **Days from germination to flowering (quadratic)** | **$F_{(1,116)}$ = 18.528 p<0.0001** | **$F_{(1,116)}$ = 7.727 p=0.006** | -- | -- | -- |
| **Initial stem diameter** | $F_{(1,116)}$ = 0.892 p=0.347 | $F_{(1,116)}$ = 0.609 p=0.437 | $F_{(1,117)}$ = 1.046 p=0.309 | $F_{(1,121)}$ = 2.456 p=0.120 | **$F_{(1,119)}$ = 5.243 p=0.024** |
| **Resource availability** | $F_{(1,116)}$ = 3.036 p=0.084 | $F_{(1,116)}$ = 1.686 p=0.197 | $F_{(1,117)}$ = 1.036 p=0.311 | $F_{(1,121)}$ = 3.282 p=0.073 | $F_{(1,119)}$ = 2.752 p=0.010 |
| **Pollination treatment** | $F_{(1,6)}$ = 0.484 p=0.513 | $F_{(1,6)}$ = 0.676 p=0.442 | $F_{(1,6)}$ = 0.153 p=0.709 | $F_{(1,6)}$ = 2.115 p=0.196 | **$F_{(1,6)}$ = 7.639 p=0.033** |
| **Resource availability x pollination treatment** | $F_{(1,116)}$ = 0.030 p=0.863 | $F_{(1,116)}$ = 0.619 p=0.433 | $F_{(1,117)}$ = 0.005 p=0.943 | $F_{(1,121)}$ = 0.000 p=0.993 | $F_{(1,119)}$ = 0.814 p=0.369 |

### FIGURE CAPTIONS

**Figure 1.** Beanplots demonstrating the association between average early anther length and the resource availability (pot size) treatment. Thin lines represent a one-dimensional scatter plot while thick lines represent the average anther length within each treatment.

**Figure 2.** The associations between (A) stem diameter at the onset of reproduction and flowering onset date, (B) average early anther length and stem diameter at the onset of reproduction, (C) average early flower diameter and stem diameter at the onset of reproduction, (D) average early ovary length and flowering onset date, and (E) early sex allocation (measured as the ratio of anther-ovary lengths in the early flower buds) and flowering onset date. Trend lines and 95% confidence intervals were extracted from the results of generalized linear mixed models that included all other explanatory variables and incorporated tray as a random effect, as shown in Table 1.

**Figure 3.** Beanplots demonstrating the association between the change in flower diameter between flower bud collection times and the pollination treatment. Thin lines represent a one-dimensional scatter plot, thick lines represent the average anther length within each treatment, and the dotted horizontal line at y=0 reflects no plasticity in flower size.

**Figure 4.** Relationships between flowering onset date and the change in (A) anther length, (B) ovary length, and (C) the anther-ovary length ratio between flower bud collection times and the relationship between stem diameter at the onset of flowering and the change in flower size between flower bud collection times. The horizontal dotted, grey lines at y=0 reflect no plasticity in ovary or anther length (or their ratio) over time. Trend lines and 95% confidence intervals were drawn from the results of generalized linear mixed models that included all other explanatory variables and incorporated tray as a random effect, as shown in Table 2.

**FIGURE 1**

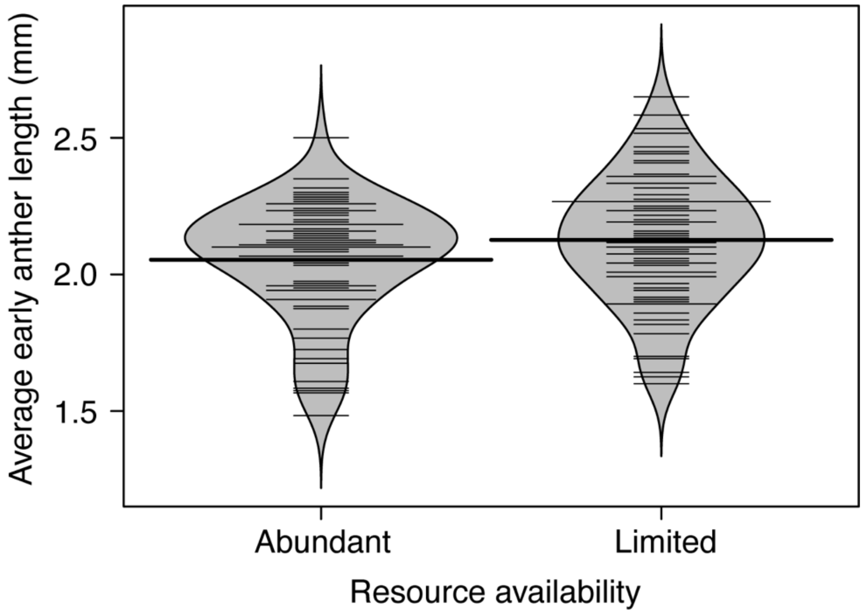

Average early anther length (mm)
2.5
2.0
1.5
Abundant
Limited
Resource availability

**FIGURE 2**

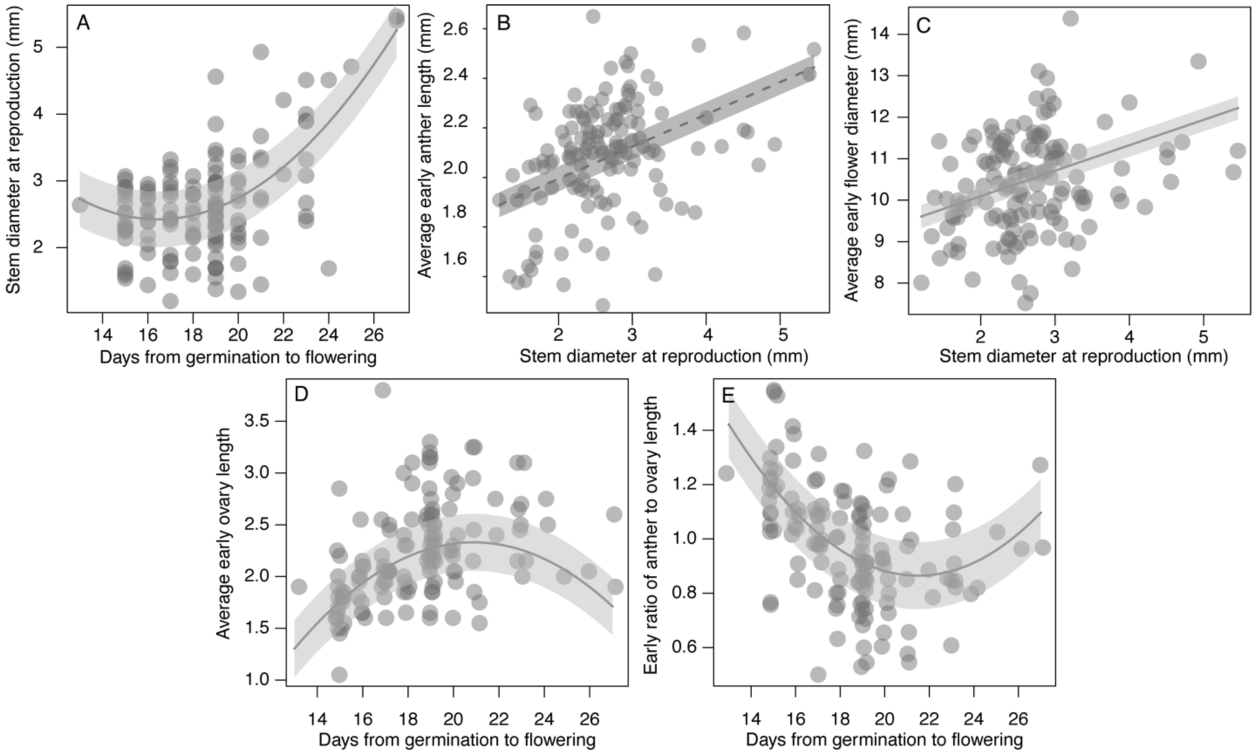


**FIGURE 3**

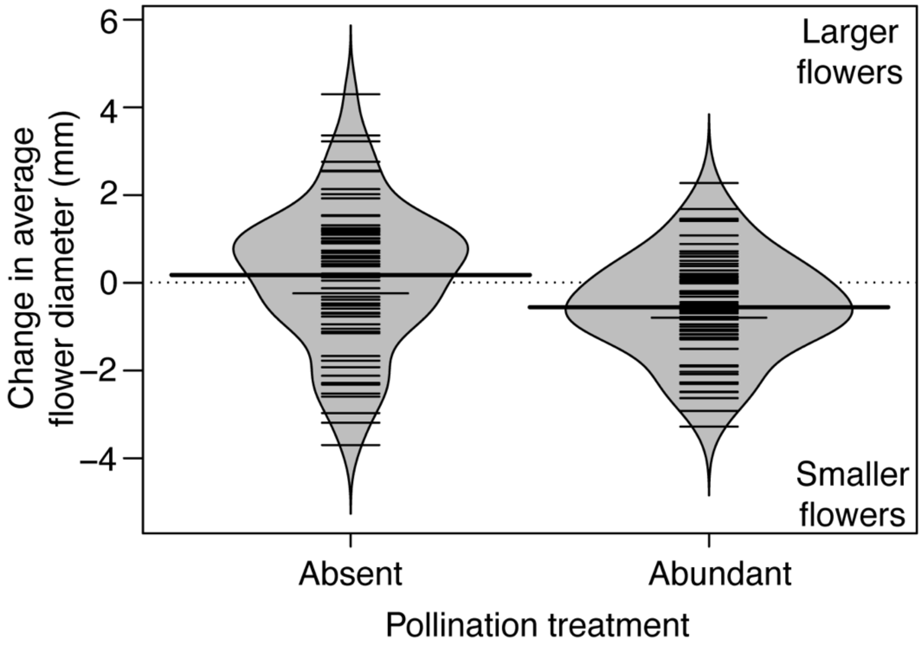

### FIGURE 4

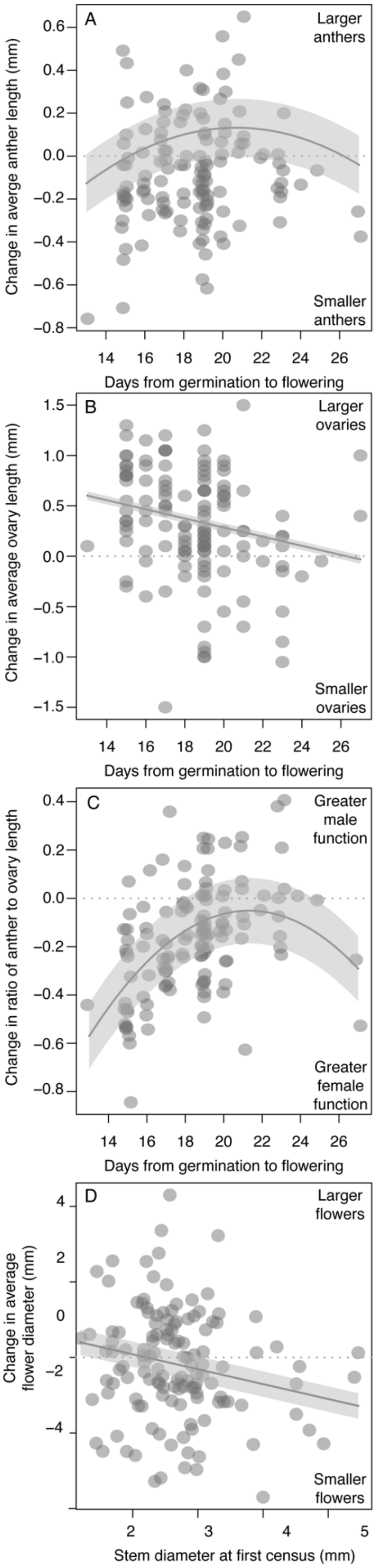
A
Larger anthers
Smaller anthers
Change in averge anther length (mm)
Days from germination to flowering
B
Larger ovaries
Smaller ovaries
Change in average ovary length (mm)
Days from germination to flowering
C
Greater male function
Greater female function
Change in ratio of anther to ovary length
Days from germination to flowering
D
Larger flowers
Smaller flowers
Change in average flower diameter (mm)
Stem diameter at first census (mm)
